# Antibiotic-Mediated Plasmonic Resonance on a Novel Nanopillar Metasurface Array

**DOI:** 10.1101/2024.10.11.617894

**Authors:** Jacob Waitkus, JaeWoo Park, Theodore Ndukaife, Sui Yang, Ke Du

## Abstract

This study demonstrates a silicon nanopillar metasurface coupled with localized surface plasmon resonance (LSPR) mediated by the presence of cephalexin antibiotics in solution for biosensing applications. A facile fabrication process was developed to create the metasurface on silicon wafers with a unique resonance signature. The resulting metasurface consists of periodic nanopillars approximately 180 nm in diameter, 210 nm deep, and with a controlled edge-to-edge separation of 200 nm. These dimensions were chosen based on a finite element method simulation that was used to investigate the ideal parameters to produce the desired resonance effect in the metasurface reflection spectra. Optimization of the nanopillar surface properties and the sidewall angle allowed for replication of the simulations. This metasurface was coupled with BSA-coated gold nanospheres (BSANS) to mediate the redshift of peak resonance wavelength values, occurring only in the presence of the antibiotic linker. The device fabricated herein exhibits a significant 22 nm wavelength shift resulting from changes to the local refractive index in the presence of the BSANS-antibiotic coupling. Further enhancement of the binding events is promoted by the LSPR hot spots formed between the nanoparticles and the metasurface allowing for sensitive and real-time detection.

## 1. Introduction

Overuse and misuse of common antibiotics has promoted antimicrobial resistance (AMR) to become a forefront issue for healthcare professionals.^1,2^ Popular medicines such as penicillin are often prescribed as a generic remedy when more specialized antibiotics would be appropriate for treatment.^3,4^ This fallacy has allowed bacterial mutations to survive and reproduce, causing new resistances and ineffective treatments. In sepsis patients, rapid and potent medication is required to combat the deadly ailment often caused by bacterial infections.^5–7^ The need for identifying AMR in these instances is thus critical. Blood work is commonly performed to identify sepsis in patients, but this detection method is not specific for the identification of AMR.^8^ To identify and monitor AMR, current techniques often involve agar dilutions and next generation sequencing (NGS). Agar dilutions are considered a gold standard for this field, cultivating multiple plates of bacteria colonies that have been exposed to varying concentrations of an antibiotic solution.^9^ The survival rate of the colonies can be recorded to determine the minimum inhibitory concentration (MIC) necessary to rid the sample of infection for a given species of bacteria. However, this process is time-consuming, requires trained professionals to operate, and is singleplexing. The NGS method allows for complete mapping of the genome from extracted DNA samples to form a library of the genetic determinants.^10,11^ This allows for easy identification of mutations in a bacteria sample that could promote AMR. However, operating and performing NGS takes specialized equipment and larger time constraints to properly establish the genetic database which is not suitable for the timely detection of AMR in sepsis patients. Comparatively, miniaturized biosensing devices may offer real-time detection and simplistic operation to promote widespread use and aid for treating afflictions.

Recent advancements in biosensing have seen the implementation of metasurface-based devices that can improve upon common techniques. This is achieved through manipulation of electromagnetic waves by the unique organization of nanoscale features smaller than the incident light wavelength.^12,13^ Through blocking or absorbing photons, altering the chirality of the wave, or enhancing the intensity of the incoming light, metasurfaces are able to be customized for various applications.^14,15^ Popular devices are able to create negative refractive index values or promote high quality factors to boost the surface sensitivity to changes in their surrounding media and improve detection limits.^16,17^ Recent research has seen a multitude of metasurfaces be applied to the biosensing field for the detection of DNA/RNA,^17^ protein structures,^18^ and antibodies.^19^ These miniaturized devices have been developed to manipulate THz waves or optical lasers for identifying viral or bacterial targets based on the surface orientation, geometry, and periodicity of the nanoscale features for the detection of weak or dilute signals.^17,19^

In addition, localized surface plasmon resonance (LSPR) detection can be facilitated through metallic nanoparticles or clusters, which promotes the collective oscillation of the conductive electron clouds of the nanomaterials to produce hot spot regions where incident photons undergo intensity enhancement.^20^ These hot spots form in the spacings between particles as the nanoparticles align their dipoles to work in unison and enhance their electron fields.^20,21^ This effect is observed only in close proximity to the particle surface as the enhancement factor exponentially diminishes with distance from the material.^21,22^ Functionalizing the surface of these materials creates differences in the local refractive index, producing different LSPR peak locations for the clusters. Our previous work has demonstrated this with LSPR-based detection of viral RNA via gold nanoparticles (AuNP) functionalized with single-stranded DNA (ssDNA).^23^ Further, coinage metals are typically employed to achieve LSPR enhancement as their collective oscillations occur within the visible spectrum or NIR region, allowing for more readily detectable changes at the particle surface and monitoring of reactions.^24^

In this work, we demonstrate a novel biosensing device by combining the metasurface with LSPR enhancement, mediated by cephalexin molecules. The silicon nanopillar metasurface was designed and modelled to produce a novel reflectance signature different than planar silicon materials, with a narrow full width at half maximum (FWHM) value below 10 nm for the specific peak of interest around 400 nm.The parameters of the nanopillars were investigated to control the location of the reflectance peak observed in the visible light region. AuNPs were coupled with an antibiotic linker that would preferentially bind to the functionalized nanopillar features. Combining LSPR and metasurface techniques, the development of an antibiotic-mediated detection assay was demonstrated, enabling a 22 nm wavelength shift resulting from changes to the local refractive index in the presence of the BSA-coated gold nanospheres (BSANS)-antibiotic coupling.

**Figure 1.**
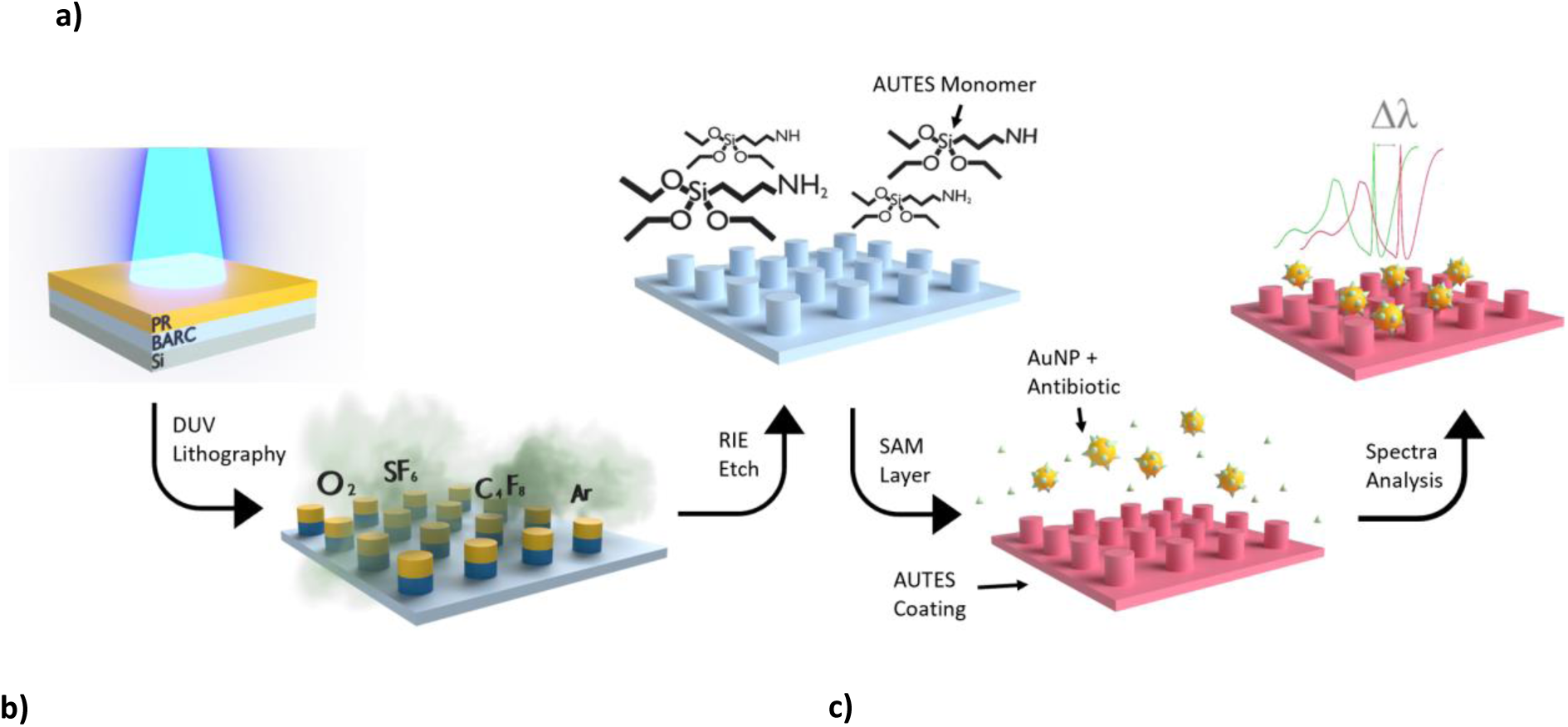
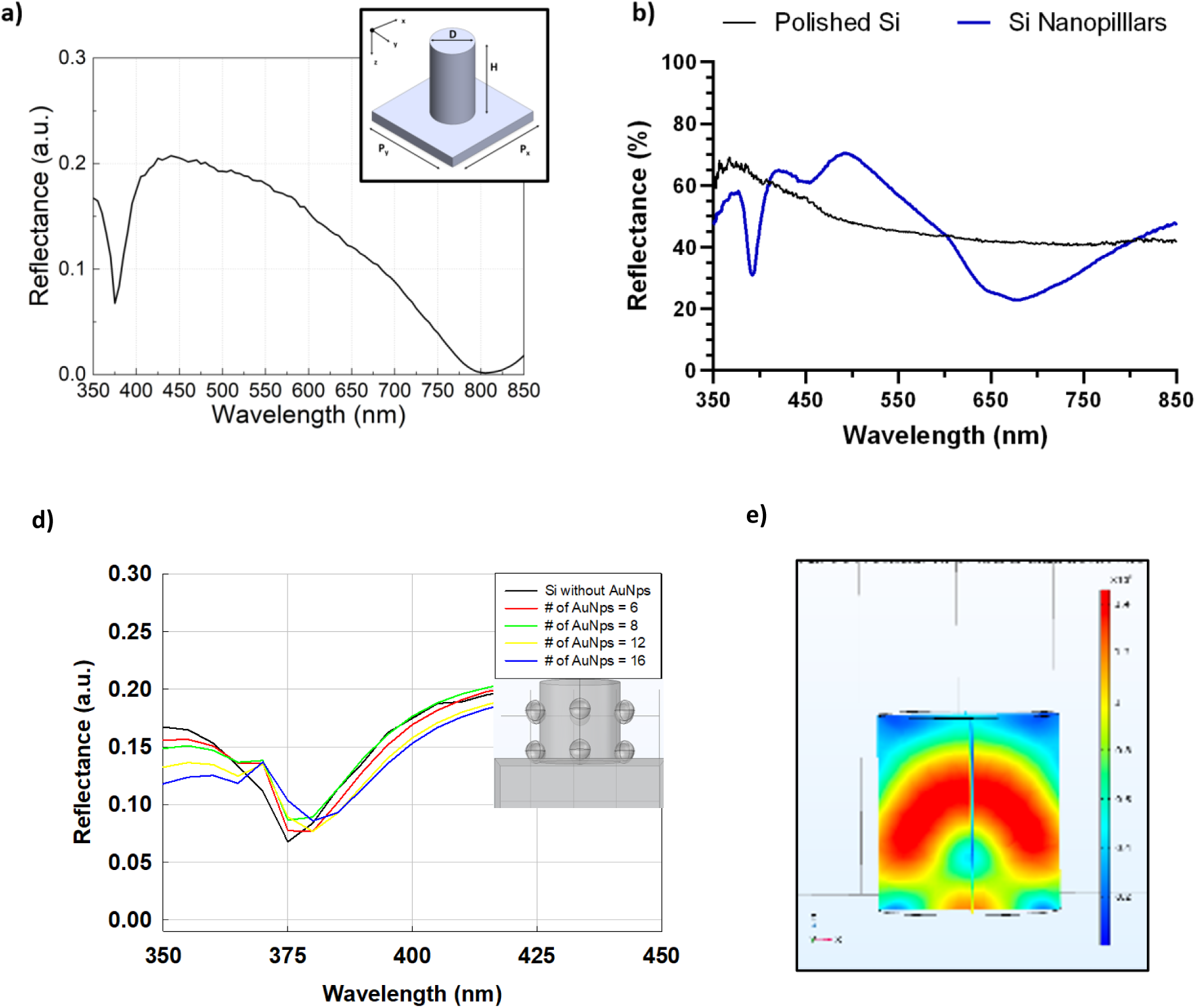
**a)** Schematic design for the fabrication and functionalization of the nanopillar metasurface for the detection of refraction index changes in the novel double-peak reflectance spectrum peak location. **b)** Simulated spectra for visible range peak reflectance. Inset highlights single pillar on substrate with critical parameters and axis labelled. **c)** Expermental reflectance spectrum for the nanopillar metasurface prior to the addition of BSANS compared to a polished silicon sample. **d)** Addition of 6-16 45nm AuNPs to the nanopillar shows a simulated redshift. e) Electron heat map for the nanopillar with greatest intensity highlighted along the sidewall.

## 2. Results and Discussion

### 2.1. Nanopillar Metasurface Array Design

The LSPR enhanced metasurface is highlighted in **Figure 1a** to demonstrate the fabrication, functionalization, and testing outlined above. Further simulations comparing the activity of the metasurface to a planar silicon wafer can be seen in **Figure 1b** and **1c**, showing two unique reflectance peaks at ∼375 nm and 670 nm, respectively. The effect of adding nanoparticles to the sidewall of the nanopillar is also simulated in **Figure 1d, 1e** for the observed redshift at the reflectance peak of interest. The presence of more nanoparticles (blue curve vs. Red curve) induces a larger redshift which can be used to detect biomarkers in the assay. By further altering the surface coating of the different LSPR particles and metasurfaces, this platform can be used to quantify enzymatic activity of resistant bacteria or mutations in bacterial DNA in sepsis patients.

### 2.2. Fabrication Optimization

The dimensions of the metasurfaces developed here pushed the resolution limit of the DUV lithography tool. To achieve the desired pattern, the system had to be optimized for the realization of the uniform PR/BARC layer across the entirety of the wafer. This was done through a focus exposure matrix (FEM) study to determine the optimal dosage to achieve the desirable nanopillars. The total thickness variation (TTV) of the silicon wafers proved to be a major contributor to the variations observed in the PR layer when performing the FEM. For an ultraflat wafer, with a TTV below 2 µm, the actual diameter of a 150 nm circle pattern varied greatly from a standard silicon test wafers as demonstrated by **Figure 2a** with the dosage exposure in mJ compared to PR diameter of the nanopillars. The importance of the wafer thickness variations can be observed optically in **Figure 2b** where the PR circle diameters and thickness produced a color change across the surface of the wafer. This is further apparent in the SEM images and accompanying reflectance spectra of the 400 nm peak in **Figure 2c and 2d** for samples from different locations on the wafer where improperly exposed PR leads to thinning that fails to protect the underlying silicon partway through the etch process. The thinner PR layers produce spike-like structures after etching which result in lower intensity peaks in the novel reflectance spectrum, more akin to that of the planar silicon substrate. The diminished effect is apparent in comparison to the nanopillar structures, where a sharper reflectance valley is observed in the visible region for the optimized lithography process.

**Figure 2.**
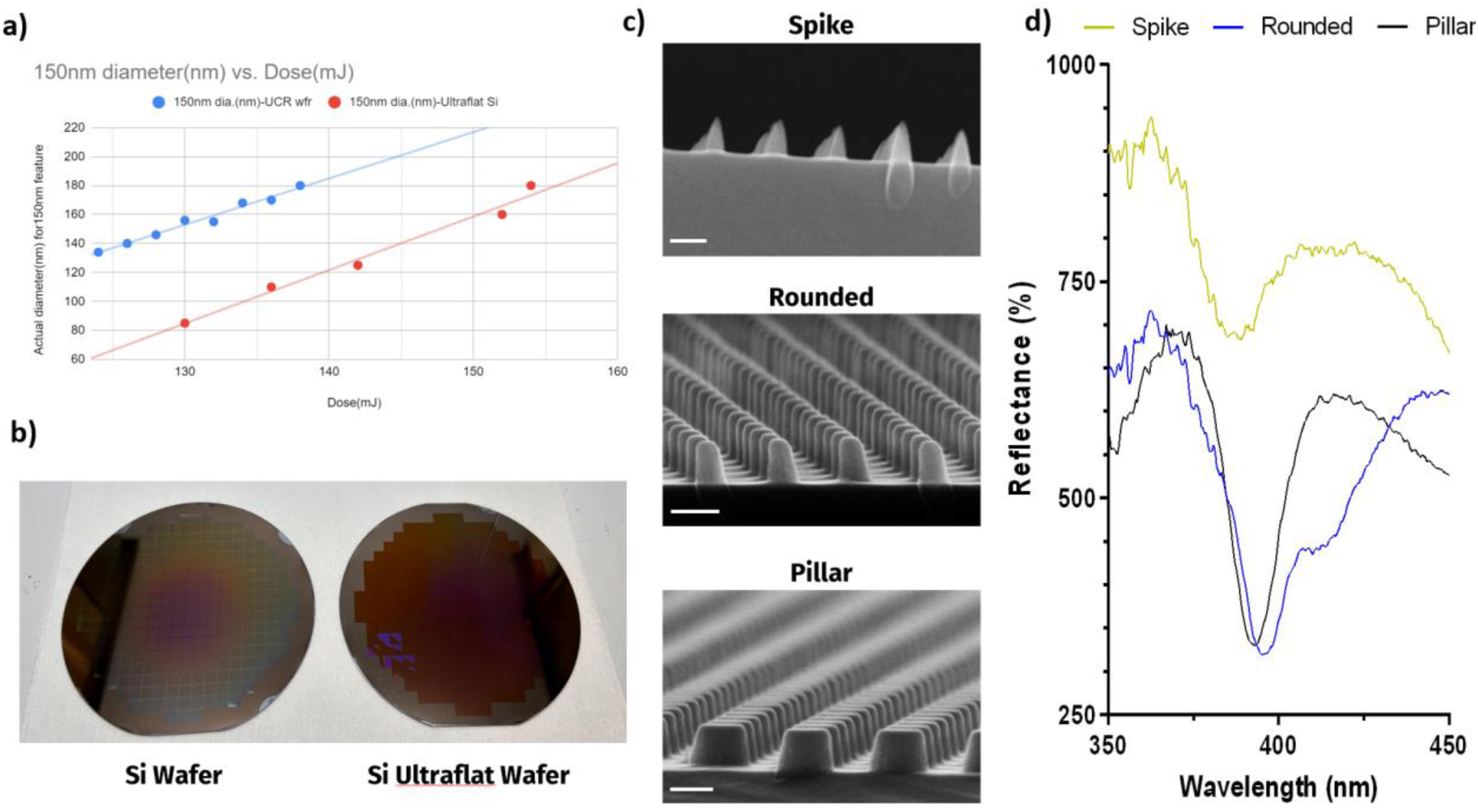
**a)** Graph for the effect of wafer thickness variation on the DUV dosage required for patterning of specific size PR circles. **b)** Optical images of wafers after DUV exposure, highlighting the PR variations on high thickness variation wafers. **c)** SEM images of three diced samples from the edge of the wafer to the center of the test wafer after etching. (Scale Bars: 200nm) **d)** Reflectance spectra for the three adjacent SEM images.

To further ensure the optimal metasurface fabrication, the sidewall angle of the nanopillar was investigated. Improving the sidewall angle helped to shrink the FWHM value of the reflectance peaks and thereby improved the sensitivity of the detection system. In the etch plasma, the ratio of the C_4_F_8_:SF_6_ was controlled between 1.22 to 5.00 as the sidewall angle was recorded by SEM images and plotted in **Figure 3a and 3b**. It was observed that above a ratio of 3.33 the sidewall angle plateaued around 84° and slowly decreased as the C_4_F_8_ ratio was increased. As the concentration of carbon in the plasma increased with respect to fluorine radicals, a greater passivation of the silicon surface occurs. This helps to promote a more anisotropic characteristic to the etch which limits undercutting of the PR/BARC layer. As the etch proceeds deeper into the silicon, the sidewalls undergo polymerization from the CF_x_ radicals in the plasma leading to the observed increase in sidewall angle. Under the RF bias, the ions in the plasma are accelerated to the surface creating a physical etch that will ignore the sidewall polymers and preferentially etch the underlying substrate, thereby further improving the sidewall angle. This effect is also controlled by the pressure value which alters the mean free path of the ion and affects the physical or chemical etch characteristic. With the implementation of the physical etch, the O_2_ gas addition aided in removal of the sidewall polymers while the Ar was crucial for diluting the F ions in the plasma and carrying away etched species. The clean surface observed in **Figure 3c** resulted in the desirable silicon nanopillars capable of producing the reflectance spectra unique to this metasurface.

**Figure 3.**
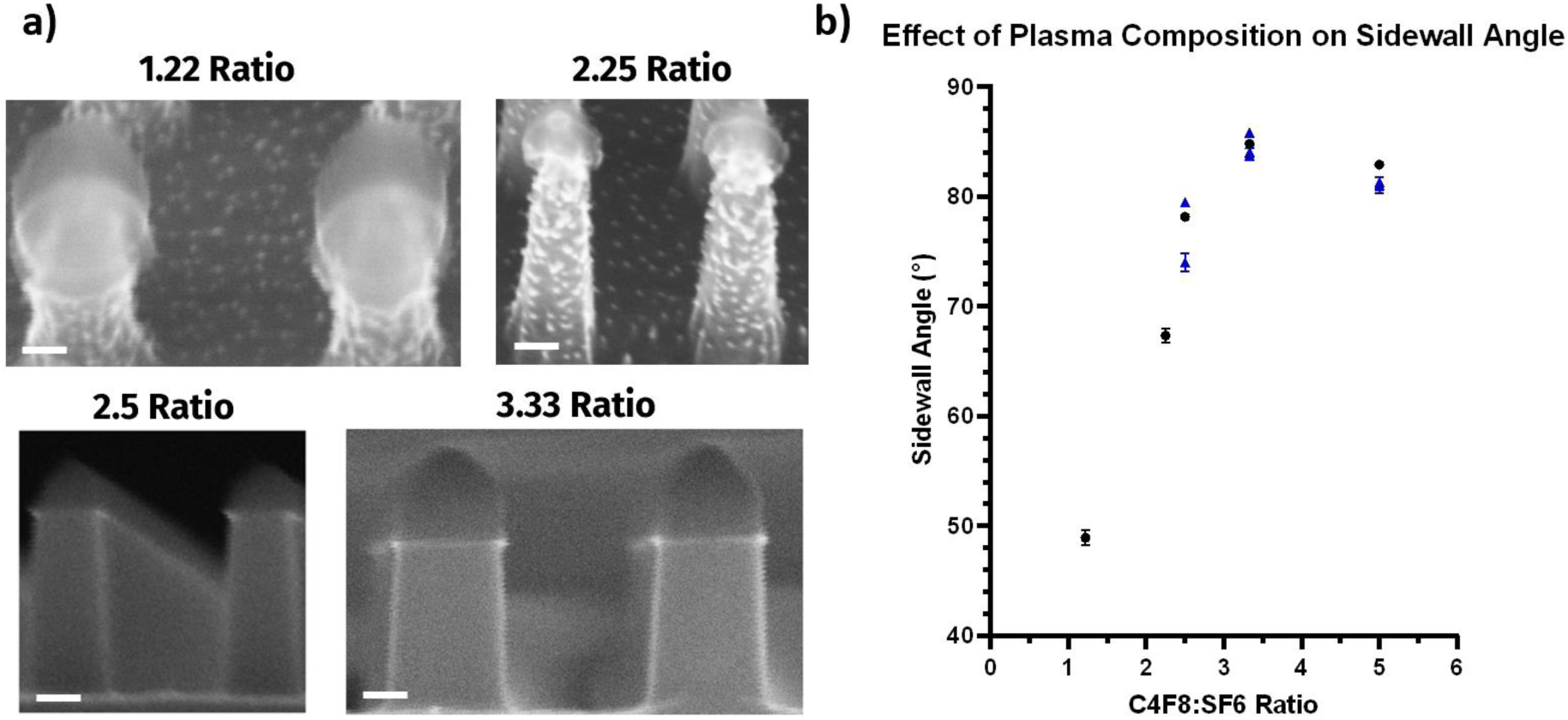
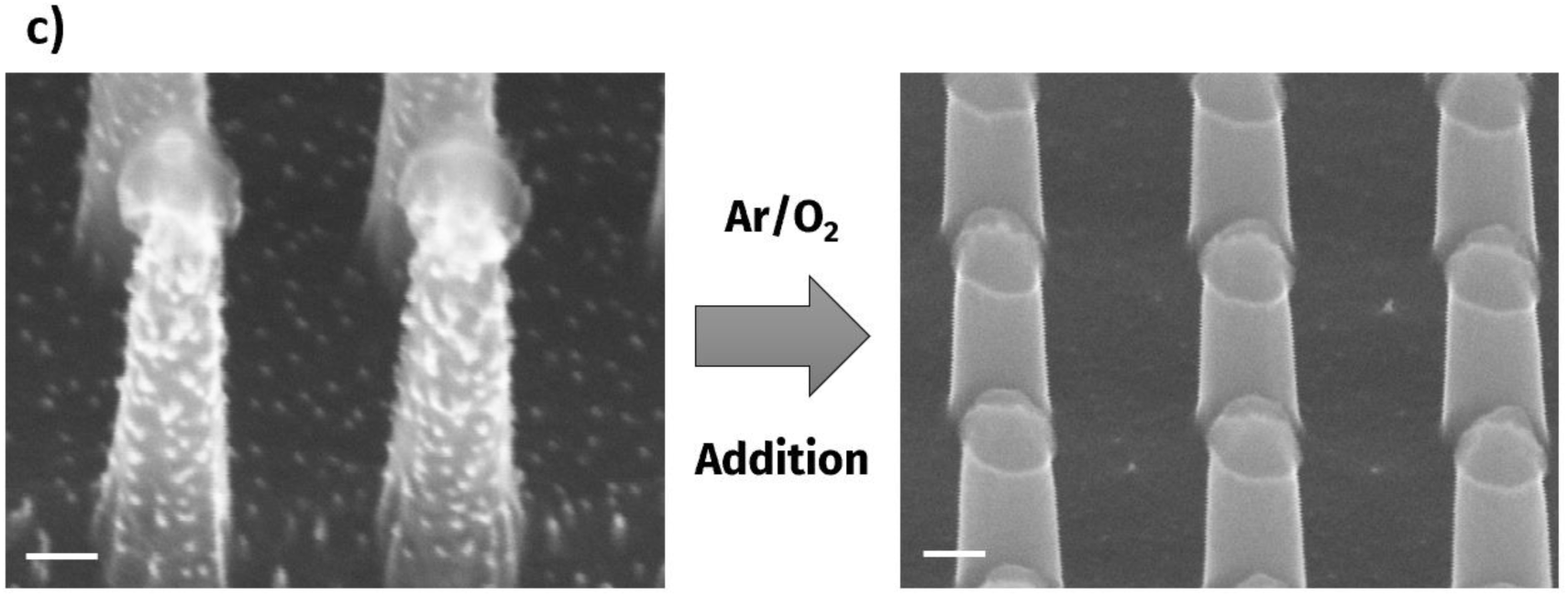
**a)** SEM images following plasma etching of Si nanopillar substrate under differing ratios of C_4_F_8_:SF_6_ gas flow rates. (Scale Bars: 100nm) **b)** Graph for the effect of C_4_F_8_:SF_6_ ratio on the sidewall angle of the Si nanopillars. **c)** SEM images highlighting the surface cleanliness following Ar/O_2_ additions to the etch plasma. (Scale Bars: 100nm)

### 2.3. LSPR Detection

To best promote the LSPR effect between the BSANS and silicon nanopillars, a coupling strategy was implemented. This consisted of a SAM layer functionalization on the nanopillars with a spacer chain present on the BSANS. The binding of these components was mediated by the presence of the cephalexin antibiotic as a linker. A multitude of different methodologies were tested to optimize the AUTES monomer layer to limit cross-linkage and multilayer formation between molecules^25,26^. The crucial factor in ideal formation was limiting the presence of water molecules on the surface. H_2_O molecules play an important role in starting the hydrolysis reaction with the siloxane groups on the AUTES chain but can lead to uncontrolled cross-linking of the monomer units if present in large quantities^27^. The ethoxy moieties on the monomer will preferentially bind to the hydroxyl groups of the native oxide layer resulting from the piranha bath previously described. Mixing the monomer solution with a toluene solvent will help to ensure monolayer formation in combination with a post-curing step to aid in alignment. The presence of the SAM layer was confirmed via contact angle measurements preceding and following each process step. With the amino terminal group coating the surface, a detectable increase in the hydrophobicity of the nanopillar surface is represented in **Figure 4a**. The AUTES coating was recorded again after 24 hours to verify the stability of the monomer when exposed to the moisture in the atmosphere. The contact angle was shown to increase from approximately 9° to over 80° via the addition of the SAM layer to the metasurface confirming the proper alignment of the terminal amino groups from the AUTES monomer on the surface of the metasurface (**Figure 4b**).

**Figure 4.**
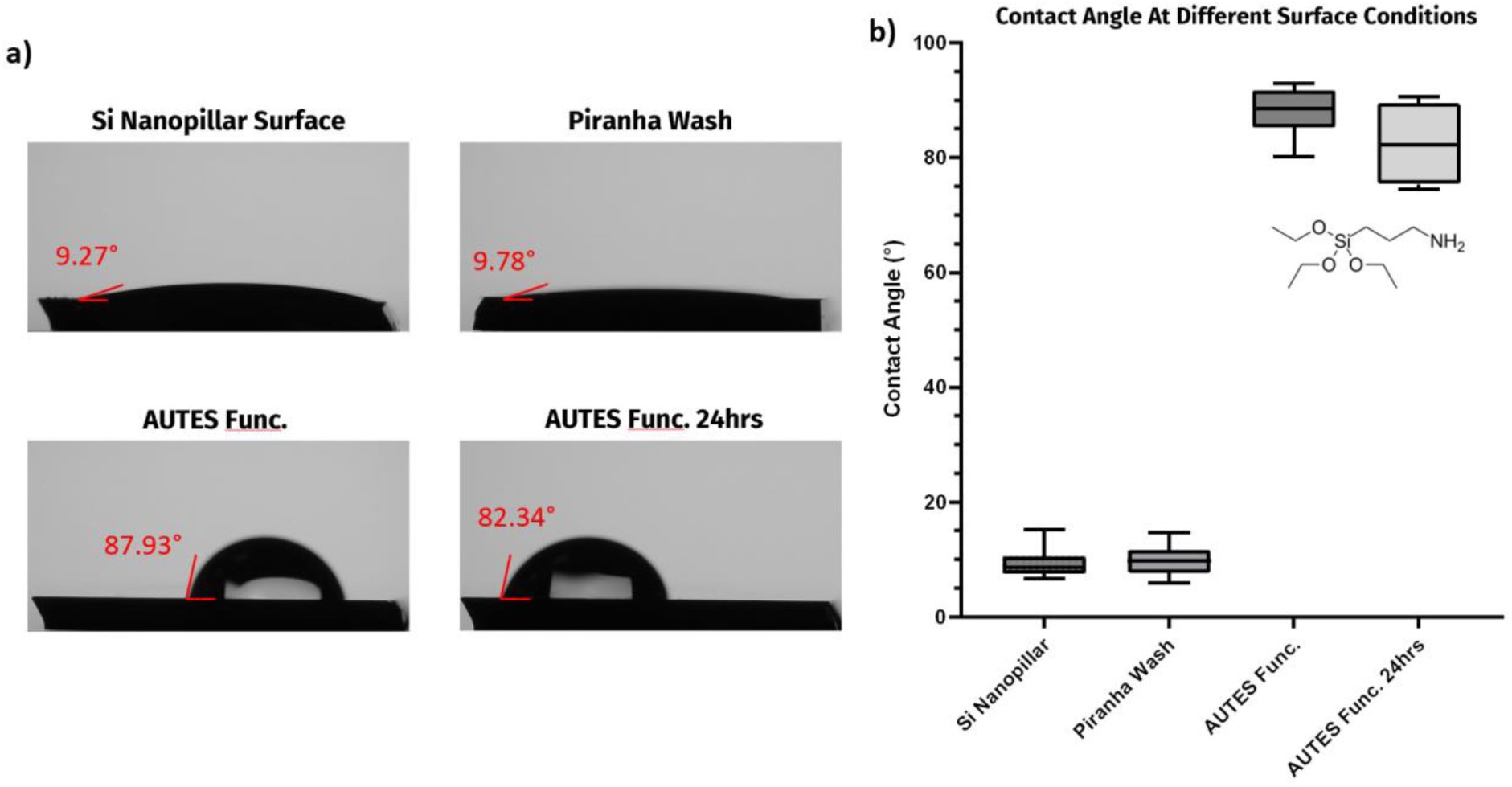
**a)** Contact angle images of a Si nanopillar surface after various process steps. **b)** Box and whisker plot for the average contact angle following the steps of Si nanopillar preparation.

The capture of the BSANS on the amino groups was promoted by the formation of a linker chain, consisting of the GLU and cephalexin antibiotic as demonstrated in **Figure 5a**. The formyl groups on opposing ends of the GLU spacer bind to the terminal amino group of lysine chains found within the BSA binding domains, and the amino group located within the cephalexin molecule^28,29^. The optimum diameter for the gold core was investigated at different linker component concentrations in **Figure 5b** to maximize the LSPR effect. For 15 nm BSANS particles, a stronger redshift was observed in the peak located at 400 nm for the reflectance spectrum when compared to the 7 nm and 45 nm particles. The SEM images in **Figure 5d** comparing the different nanoparticles sizes shows the greater coverage of the nanopillar sidewalls with the smaller particles when compared to that of the 45 nm BSANS.

Figure 5c graphs the redshift in reflectance peak wavelength under differing ratios for the GLU and cephalexin molecules. The optimal condition was observed to be at 10,000 times greater GLU molecules than BSA binding sites and 100 times greater cephalexin compounds compared to the number of BSA proteins, where the wavelength shift detected was over 20 nm. The oversaturation of the linkers allowed for an increased functionalization of the BSANS binding locations to improve their capture on the silicon features. As the presence of the GLU decreased in solution, the detectable shift decreased to a minimum redshift around 3 nm. In the absence of GLU spacers, the effect of cephalexin concentration exhibited a limited effect where all samples produced a minute redshift between 3 and 5 nm (blue bars). This result aligns with the protein envelopment concept as without a spacer the gold core would struggle to combine with the silicon metasurface^30,31^, only producing occasional non-specific binding resulting in the small-scale changes detected here.

**Figure 5.**
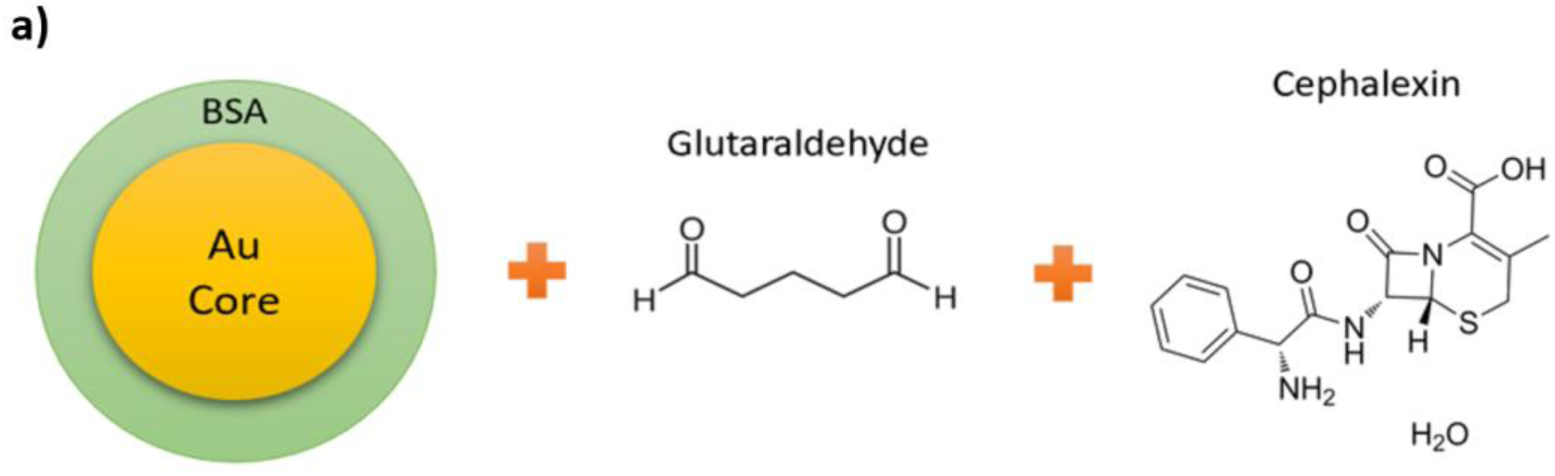
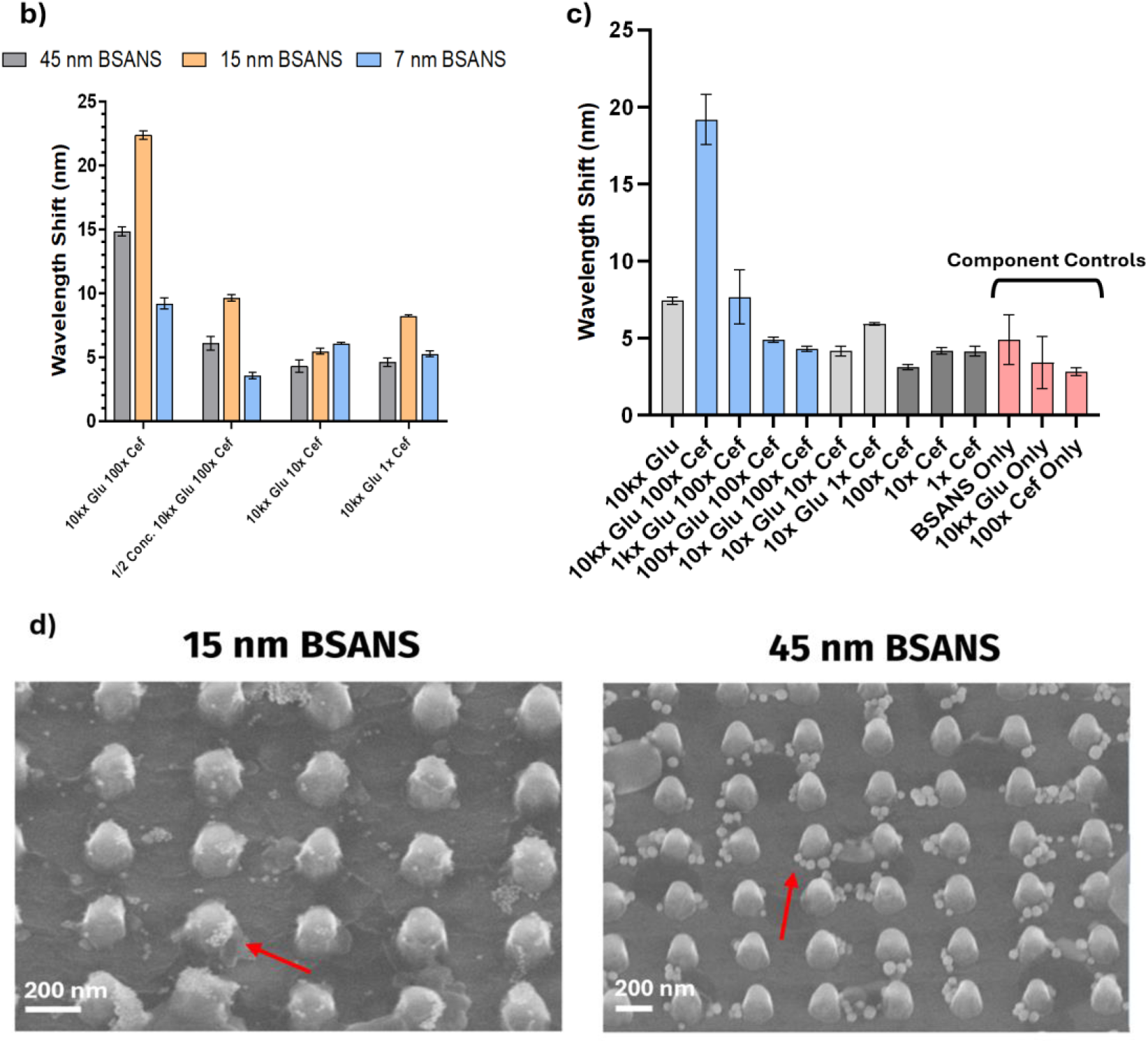
**a)** Model of the BSA-coated AuNP (BSANS) linker formation to bind to the functionalized Si nanopillars. **b)** Effect of the Au core on the magnitude of redshift detected under various linker concentrations. **c)** Study on the effects of altering the glutaraldehyde and cephalexin concentrations for the maximum BSANS capture. d) SEM images for the 15nm and 45nm BSANS particles, showing the coverage of the pillars under identical conditions. (Scale Bars: 200nm)

## 3. Conclusion

The silicon-based nanopillars fabricated and tested in this work establish a LSPR and metasurface device capable of sensitive detection of AuNPs in the presence of a target antibiotic. The manipulation of incident electromagnetic waves was achieved by precise control over the nanoscale features to produce the reflectance spectrum with a narrow FWHM value and accompanying sensitive secondary peaks. Through a simple production process, the geometry of the silicon nanopillars was optimized to best promote this simulated effect. Control over the sidewall angle of the pillar and limiting the undercutting of the PR/BARC layer proved crucial in production of the novel metasurface. Through coupling with LSPR enhancement, the capture of BSANS on the surface provide a mechanism for detection of the presence of cephalexin in solution. Optimization of the BSANS particle diameter highlighted the importance of complete nanopillar coverage. The larger BSANS preferentially bound to the crux between the nanopillar and the planar substrate. Based on the electron heat map in our simulations, the electron interactions were strongest along the nanopillar sidewall. This region would produce a stronger LSPR effect in the form of nanoparticle hot spots and evanescent waves in close proximity to the Si nanopillar surface. Larger metallic nanoparticles are known to have an enhanced LSPR effect due to their greater electron cloud mobility and increased hot spot strength, but the formation of higher order clusters and greater nanopillar coverage by the 7 nm and 15 nm BSANS helps them overcome their weaker expected LSPR property through stronger interactions with the nanopillar substrate. The 15 nm gold core would still produce a stronger effect than that of the 7 nm BSANS, while being easier to observe in a SEM and detect so they were chosen as the optimal condition.

Implementation of the GLU molecule as a space was to improve the sensitivity of the testing assay. Without the presence of the GLU molecule, the protein binding domains envelop the cephalexin antibiotic. This encompassing creates steric hinderance that limits the ability for the antibiotic binding sites to couple with the AUTES amino group on the silicon nanopillars thereby decreasing the overall LSPR effect with the metasurface. In future experiments, this complete coverage of the bound antibiotic would limit the devices applicability for testing of AMR. The BSA protein would prevent capture of the intact antibiotic molecules from binding to penicillin binding proteins and inactivating them by blocking the terminal antibiotic groups. Future work on this metasurface would allow for the detection of AMR in the form of enzymatic activity or genetic mutations by varying the linker chain and surface functionalization of the silicon to the appropriate capture materials.^32,33^

Our unique metasurface structure provides two distinct signature peaks which are over 300 nm apart from each other. In the future, these additonal peaks can be developed as internal controls for multiplexing sensing.^34^ In addition, patterned by DUV lithography, numerous chips can be used in parallel, which is ideal for the detection of multiple analytes in a patient sample. The same sample can potentially be used for multiplexed detection as well through careful customization of the nanoparticles to produce shifts on separate regions of the metasurface reflectance spectrum.^35^ Fabrication of these metasurfaces can be done on flexible surfaces as well, allowing for coating of fiber optics or other like materials to improve their inherent properties or to promote detection^36,37^. Expanding the applicability of the nanopillar metasurface promotes the viability of the detection strategy for various analytes. The realization of this device demonstrates the advantage of real-time and sensitive detection strategies capable of multiplex detection.

## 4. Experimental Section/Methods

### 4.1. Nanopillar Fabrication

For the realization of the nanopillar metasurface detection assay, a facile fabrication process was developed requiring only nanolithography and dry etch processing, followed by the functionalization of the metasurface to integrate the LSPR-based detection assay. The patterning was performed using a deep-UV (DUV) photolithography tool (ASML 5500 stepper) with an exposure area confined to the overlap of a 31 mm diameter circle and a 22 mm x 27 mm rectangle. The mask used herein contained nine various nanopillar patterns on a 6” quartz wafer (6250E grade; Didgitat-Toppan) allowing for multiple different dimensions to be patterned with ease. The metasurface lithography was achieved via exposure of a 248 nm KrF laser from the DUV tool onto a UVN30 negative photoresist (PR) and a DUV-42P-6 bottom antireflective coating (BARC) that was spun onto a 4”/100 mm ultraflat silicon wafer (Ultrasil) at a thickness of 500 nm and 80 nm respectively^38^. The unexposed PR layer was removed via an AZ300MIF developer wash. To remove the BARC layer from the unexposed regions, an O_2_ plasma etch was performed in an FL-ICP (Plasma-Therm 770 SLR) tool under 20 sccm of O_2_ at 10 mTorr with a 100 W RF bias for 1 min^39^.

The nanopillar formation was finalized by a reactive ion etching (RIE) process to produce the desirable nanopillar height and sidewall conditions. Prior to the etch, the patterned wafers were cleaved by diamond-tipped scribe pens into approximately 2 cm^2^ samples and cleaned under an air jet. This was done to gently remove debris that resulted from the cleavage without altering the remaining PR/BARC pattern. The cleaved samples were set on a 4” silicon carrier wafer and inserted into an atomic layer etching (ALE) tool (PlasmaPro 100 ALE System) that was used in RIE mode. The plasma was comprised of C_4_F_8_, Ar, SF_6_, and O_2_, at 50, 30, 15, and 2 sccm respectively. These gases were ignited and controlled by an RF bias at 50 W and ICP power set at 1200 W under 10 mTorr of pressure. This process was run for 2 mins on the empty ALE chamber prior to etching the samples to condition the chamber for production. The etch time was adjusted depending on nanopillar height, where a 51 s was used to produce 210 nm deep nanopillars. To remove the remaining PR/BARC layers, an O_2_/Ar strip was performed in the same chamber with 50 sccm of O_2_, 10 sccm of Ar, at 20 mTorr pressure, a bias of 75 W, and an ICP of 2,000 W for 5 mins. Between the etch and the strip step, the samples were removed, and the strip step was run to ensure the chamber was properly conditioned to limit any further silicon etching by remaining etchants from the previous step. This allowed for the final realization of nanopillar metasurface.

### 4.2. Testing Apparatus

Reflectance spectrums were recorded via the Ocean Insight Flame Miniature Spectrometer (**Figure S1**). A tungsten halogen lamp (HL-2000 Ocean Insight) was directed to the surface through a collection of fiber optic cables. The cable bundle consisted of six emission fiber optics surrounding the detector probe (QR400-7-SR-BX EOS-A695605 OceanInsight). Silicon nanopillar samples were tested using a Spectralon (Labsphere) background surface standard, which is reflective over 99% for the visible wavelength region. Contact angle measurements were performed using a contact angle goniometer from Ossila. A Canny edge detection algorithm was used to record the contact angle of 5 µL water droplets on the surface of the metasurface. Nanopillar samples were air jet dried to preserve the surface condition.

### 4.3. Modelling

To characterize the silicon metasurface performance, 3D electromagnetic simulations using finite element methods were performed within COSMOL Multiphysics. This was done to design the optimal periodicity, diameters, and nanopillar height to achieve the metasurface reflectance spectrum on the polarization insensitive metasurface without sacrificing the resonance linewidth/FWHM. For the COSMOL simulations, port boundary conditions were applied along the direction of propagation of incident radiation with periodic boundary conditions along the x and y axes. Perfectly matched layer absorbing boundary conditions were also applied in the z direction. Refractive index changes were also simulated in the presence of bound 45 nm gold nanoparticles to observe the resulting redshift and increasing intensity from the LSPR effect across the complete visible spectrum when compared to the experimental spectrum of the metasurface and a planar silicon sample.

### 4.4. Antibiotic Detection Assay

Towards the combination of LSPR effects with metasurface, an assay was developed incorporating antibiotics as the linker system between BSANS and the silicon nanopillars. Three sizes of BSANS were acquired from Luna Nanotech at stock concentrations of 4.9 nM for the 45 nm particles, 0.136 µM for the 15 nm particles, and 1.56 µM for the 7 nm diameter BSANS. All three samples were stored at 4°C. A glutaraldehyde (GLU) molecule was implemented as a spacer between the BSA coating and the cephalexin antibiotic. The GLU was suspended in aqueous media at 2.5% (Electron Microscopy Sciences) and stored at 4°C. The cephalexin was purchased from Thermo Fisher Scientific as a monohydrate powder and stored at 4°C as well. For the self-assembled monolayer (SAM) on the nanopillars, an 11-aminoundecyltriethoxysilane (AUTES) monomer was purchased from Gelest, Inc.

The silicon nanopillar surface was functionalized with a SAM layer following a pre-treatment procedure. The cleaved samples were sonicated in acetone and IPA separately for 5 mins before being dried via air jet. The samples were then submerged in a piranha solution for 30 mins at room temperature. This bath was prepared at a 3:1 ratio of H_2_SO_4_:H_2_O_2_ by volume. Immediately following the piranha solution, the samples were washed in copious amounts of DI water and dried under N_2_ gas. Here, the piranha wash was used to remove any organic contaminants and to form a uniform and thin silicon oxide layer which is preferable for the capture of the AUTES monomer. For functionalization of the silicon with the monomer, the samples were washed in toluene three times to limit water presence on the surface. The AUTES monomer was dissolved in toluene to 2% at 90°C for 2 hrs. The ideal solution condition was tested and shown in **Figure S2**. These samples were then washed in toluene three times again to remove any unbound molecules from the surface. The final devices were baked at 110°C for 30 mins to ensure the monomer alignment was ideal and to remove any remaining moisture.

The BSANS linker chain was formed by first mixing 20 µL of stock BSANS with 200 µL of GLU for the 7 nm and 15 nm particles, while 20 µL of the 45 nm particles were mixed with 20 µL of GLU. This was done to ensure that the ratio of GLU molecules to the number of BSA lysine chains was consistent at 10,000x. The solution was mixed for 5 mins before introducing the cephalexin antibiotic, followed by another 30 mins of mixing at room temperature. The antibiotic was dissolved in PBS 1x (pH 7.4) to a final concentration of ∼800 µM for the 7 nm and 15 nm particles, and ∼80 µM for the 45 nm particles. Finally, the device was washed in PBS 1x three times, and air jet dried to remove any weakly associated or unbound nanoparticle chains before spectral testing.

## Supporting Information

Supporting Information is available from the Wiley Online Library or from the author.

## Acknowledgements

This work is supported by the National Institutes of Health (No. R35GM 142763) and the National Institute of Food and Agriculture (2024-67021-42833) The lithography work was performed with the help of the University of California, Santa Barbara. Gopikrishnan Meena performed the exposures and process development, Demis D. John designed the process originally reported by Brian Thibeault and provided analysis and direction.

Received: ((will be filled in by the editorial staff))

Revised: ((will be filled in by the editorial staff))

Published online: ((will be filled in by the editorial staff))

## References

1. Swami, O. C. Strategies to Combat Antimicrobial Resistance. JCDR (2014) doi:10.7860/JCDR/2014/8925.4529.

2. Global burden of antimicrobial resistance: essential pieces of a global puzzle – Authors’ reply. The Lancet 399, 2349–2350 (2022).

3. 3. Antibiotics: Are you misusing them? Mayo Clinic https://www.mayoclinic.org/healthy-lifestyle/consumer-health/in-depth/antibiotics/art-20045720.

4. Batista, A. D. et al. Antibiotic Dispensation without a Prescription Worldwide: A Systematic Review. Antibiotics 9, 786 (2020).

5. Sepsis. Medicine 45, 649–653 (2017).

6. Mayr, F. B., Yende, S. & Angus, D. C. Epidemiology of severe sepsis. Virulence 5, 4–11 (2014).

7. Oeschger, T., McCloskey, D., Kopparthy, V., Singh, A. & Erickson, D. Point of care technologies for sepsis diagnosis and treatment. Lab on a Chip 19, 728–737 (2019).

8. Sheldon, I. M. Detection of Pathogens in Blood for Diagnosis of Sepsis and Beyond. eBioMedicine 9, 13–14 (2016).

9. Methods for Dilution Antimicrobial Susceptibility Tests for Bacteria That Grow Aerobically: M07-A10 ; Approved Standard. (Committee for Clinical Laboratory Standards, Wayne, PA, 2015).

10. Diagnosis of Sepsis with Cell-free DNA by Next-Generation Sequencing Technology in ICU Patients. Archives of Medical Research 47, 365–371 (2016).

11. Grumaz, S. et al. Next-generation sequencing diagnostics of bacteremia in septic patients. Genome Med 8, 73 (2016).

12. Zhou, T. et al. A Metasurface Plasmonic Analysis Platform Combined with Gold Nanoparticles for Ultrasensitive Quantitative Detection of Small Molecules. Biosensors 13, 681 (2023).

13. Li, A., Singh, S. & Sievenpiper, D. Metasurfaces and their applications. Nanophotonics 7, 989–1011 (2018).

14. Bukhari, S. S., Vardaxoglou, J. (Yiannis) & Whittow, W. A Metasurfaces Review: Definitions and Applications. Applied Sciences 9, 2727 (2019).

15. Yu, N. & Capasso, F. Flat optics with designer metasurfaces. Nature Mater 13, 139–150 (2014).

16. Wang, J., Allein, F., Boechler, N., Friend, J. & Vazquez-Mena, O. Design and Fabrication of Negative-Refractive-Index Metamaterial Unit Cells for Near-Megahertz Enhanced Acoustic Transmission in Biomedical Ultrasound Applications. Phys. Rev. Appl. 15, 024025 (2021).

17. Hu, J. et al. Rapid genetic screening with high quality factor metasurfaces. Nat Commun 14, 4486 (2023).

18. High-sensitivity biosensor for identification of protein based on terahertz Fano resonance metasurfaces. Optics Communications 473, 125850 (2020).

19. 19. All-Dielectric Metasurface Fluorescence Biosensors for High-Sensitivity Antibody/Antigen Detection | ACS Nano. https://pubs.acs.org/doi/full/10.1021/acsnano.0c07722.

20. Kastner, S. et al. LSPR-Based Biosensing Enables the Detection of Antimicrobial Resistance Genes. Small 19, 2207953 (2023).

21. Tian, L. et al. Rational Approach to Plasmonic Dimers with Controlled Gap Distance, Symmetry, and Capability of Precisely Hosting Guest Molecules in Hotspot Regions. J. Am. Chem. Soc. 143, 8631–8638 (2021).

22. Wang, C. et al. Importance of Hot Spots in Gold Nanostructures on Direct Plasmon-Enhanced Electrochemistry. ACS Appl. Nano Mater. 1, 5805–5811 (2018).

23. Waitkus, J. et al. Gold Nanoparticle Enabled Localized Surface Plasmon Resonance on Unique Gold Nanomushroom Structures for On-Chip CRISPR-Cas13a Sensing. Advanced Materials Interfaces 10, 2201261 (2023).

24. LSPR-based nanobiosensors. Nano Today 4, 244–251 (2009).

25. Comparison of modified molecules 3-aminopropyltriethoxysilane and 11-aminoundecyltriethoxysilane in orientation angle and interaction with protein by sum frequency vibration spectrum and imaging ellipsometry biosensor. Thin Solid Films 769, 139738 (2023).

26. Aissaoui, N., Bergaoui, L., Landoulsi, J., Lambert, J.-F. & Boujday, S. Silane Layers on Silicon Surfaces: Mechanism of Interaction, Stability, and Influence on Protein Adsorption. Langmuir 28, 656–665 (2012).

27. Sypabekova, M., Hagemann, A., Rho, D. & Kim, S. Review: 3-Aminopropyltriethoxysilane (APTES) Deposition Methods on Oxide Surfaces in Solution and Vapor Phases for Biosensing Applications. Biosensors 13, 36 (2023).

28. Huang, B. X., Kim, H.-Y. & Dass, C. Probing three-dimensional structure of bovine serum albumin by chemical cross-linking and mass spectrometry. J Am Soc Mass Spectrom 15, 1237–1247 (2004).

29. Sandison, M. E., Cumming, S. A., Kolch, W. & Pitt, A. R. On-chip immunoprecipitation for protein purification. Lab Chip 10, 2805–2813 (2010).

30. Hamishehkar, H., Hosseini, S., Naseri, A., Safarnejad, A. & Rasoulzadeh, F. Interactions of cephalexin with bovine serum albumin: displacement reaction and molecular docking. Bioimpacts 6, 125–133 (2016).

31. Miller, L. M. et al. Antibiotic-functionalized gold nanoparticles for the detection of active β-lactamases. Nanoscale Adv. 4, 573–581 (2022).

32. Zhuang, Q. et al. Advances in the detection of β-lactamase: A review. International Journal of Biological Macromolecules 251, 126159 (2023).

33. Liu, Y. et al. One-tube RPA-CRISPR Cas12a/Cas13a rapid detection of methicillin-resistant *Staphylococcus aureus*. Analytica Chimica Acta 1278, 341757 (2023).

34. Wang, S., Liu, S., Ni, W., Wu, S. & Lu, P. Dual-wavelength highly-sensitive refractive index sensor. *Opt. Express*, OE 25, 14389–14396 (2017).

35. Pearce, A. K., Wilks, T. R., Arno, M. C. & O’Reilly, R. K. Synthesis and applications of anisotropic nanoparticles with precisely defined dimensions. Nat Rev Chem 5, 21–45 (2021).

36. Zhao, Q. et al. Optical Fiber-Integrated Metasurfaces: An Emerging Platform for Multiple Optical Applications. Nanomaterials (Basel) 12, 793 (2022).

37. Zhang, X., Cai, H., Rezaei, S. D., Rosenmann, D. & Lopez, D. A universal metasurface transfer technique for heterogeneous integration. Nanophotonics 12, 1633–1642 (2023).

38. J. Morris, N., et al. Laser desorption ionization (LDI) silicon nanopost array chips fabricated using deep UV projection lithography and deep reactive ion etching. RSC Advances 5, 72051–72057 (2015).

39. Du, K., Liu, Y., Wathuthanthri, I. & Choi, C.-H. Dual applications of free-standing holographic nanopatterns for lift-off and stencil lithography. Journal of Vacuum Science & Technology B 30, 06FF04 (2012).

