## Supporting information for "Antibiotic-Mediated Plasmonic Resonance on a Novel Nanopillar Metasurface Array"

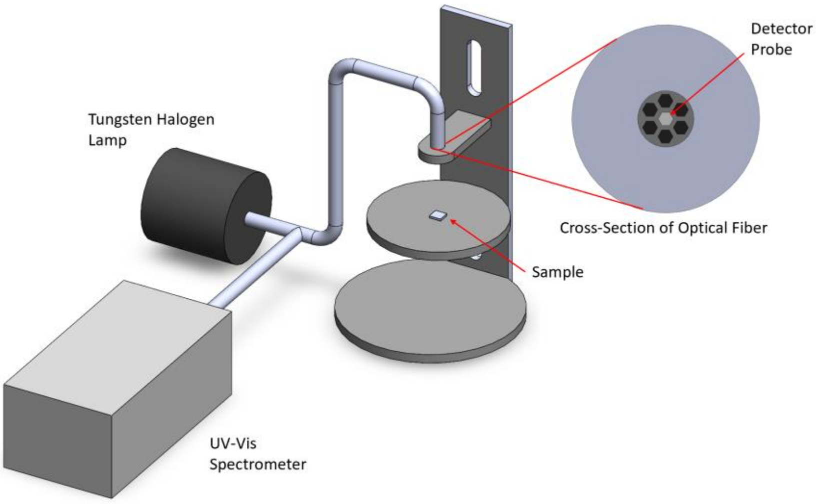


**Supplementary Figure 1.** 3D model for the reflectance spectroscopy measurements containing the tungsten halogen lamp light source, the UV-Vis spectrometer, and the Y-branched fiber optic coupling. The cross section of the optical fiber is shown in the top right consisting of six emitter probes surrounding a single detector probe.

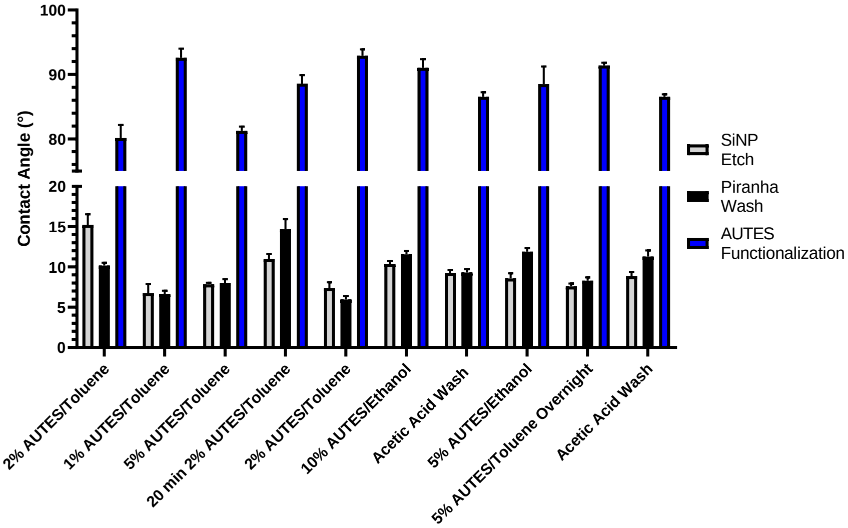


**Supplementary Figure 2.** Bar chart for different AUTES functionalization solutions. Solutions of toluene and ethanol were tested to find the optimal alignment and SAM layer formation.
